# Di-valent siRNA Mediated Silencing of MSH3 Blocks Somatic Repeat Expansion in Mouse Models of Huntington’s Disease

**DOI:** 10.1101/2022.09.06.506795

**Authors:** Daniel O’Reilly, Jillian Belgrad, Chantal Ferguson, Ashley Summers, Ellen Sapp, Cassandra McHugh, Ella Mathews, Julianna Buchwald, Socheata Ly, Dimas Echeverria Moreno, Zachary Kennedy, Vignesh Hariharan, Kathryn Monopoli, X. William Yang, Jeffery Carroll, Marian DiFiglia, Neil Aronin, Anastasia Khvorova

## Abstract

Huntington’s Disease (HD) is a severe neurodegenerative disorder caused by expansion of the CAG trinucleotide repeat tract in the huntingtin gene. Inheritance of expanded CAG repeats is needed for HD manifestation, but further somatic expansion of the repeat tract in non-dividing cells, particularly striatal neurons, hastens disease onset. Called somatic repeat expansion, this process is mediated by the mismatch repair (MMR) pathway. Among MMR components identified as modifiers of HD onset, MutS Homolog 3 (MSH3) has emerged as a potentially safe and effective target for therapeutic intervention. Here, we identify fully chemically modified short interfering RNA (siRNA) that robustly silence MSH3 *in vitro* and *in vivo*. When synthesized in a di-valent scaffold, siRNA-mediated silencing of MSH3 effectively blocked CAG repeat expansion in striatum of two HD mouse models without impacting tumor-associated microsatellite instability. Our findings establish a novel paradigm for treating patients with HD and other repeat expansion diseases.

**One Sentence Summary:** Silencing MSH3 in the CNS of two models of Huntington’s disease using di-valent siRNA blocks disease-accelerating somatic expansion of CAG repeats.

## INTRODUCTION

Huntington’s Disease (HD) is a rare autosomal dominant neurodegenerative disease that impairs cognitive and motor function, eventually leading to death (*1, 2*). Currently, no disease-modifying treatments are available (*3*). HD is caused by an expansion of the CAG repeat tract in the huntingtin gene (*HTT*), with age of disease onset being strongly driven by the number of CAG repeats (*4-6*). Individuals with ≥40 CAG repeats develop HD in their 40s whereas individuals with ≥70 repeats develop juvenile-onset HD.

CAG repeat number is inherited, but undergoes expansion over time due to somatic instability (*7*). This process, termed somatic repeat expansion, occurs preferentially in non-dividing cells with active transcription (*8*), such as neurons, and generates significant mosaicism in patient brains (*7, 9*). Somatic repeat expansion occurs when repetitive DNA (i.e., sequential CAGs) misaligns during transcription, creating a slipped loop intermediate that recruits mismatch repair (MMR) machinery to cleave the opposite (non-slipped) strand (*10-12*). The slipped loop then is used as a template to add new nucleotides that further expand the locus (*10-12*).

A recent genome-wide association study identified several MMR genes as major modifiers of both HD onset (*13*), expansion of the CAG repeat tract (*8, 13*), and clinical HD progression (*14*) suggesting this pathway as a potential therapeutic target for HD. MMR is pivotal to maintaining cellular function, repairing single base mismatches, deletions, and small and large loops to prevent genomic instability and carcinogenesis (*15, 16*). Mutations in MMR genes are associated with cancers, including those affecting the brain (*15, 16*). Thus, development of an expansion-modifying therapy for HD requires careful selection of an MMR gene target.

Among MMR gene candidates, MutS Homolog 3 (MSH3) emerges as a potentially safe and effective target for knockdown. Forming a complex with MSH2, MSH3 (MutSβ) selectively recognizes large (>3 nucleotide) DNA loops, such as those created by expanded CAG repeats, and is not involved in other pathways essential for maintenance of DNA integrity (*17*). Single nucleotide polymorphisms in the *MSH3* gene are associated with enhanced levels of CAG expansion (*13, 18*), as well as colon cancer (*15, 16*), but critically, are not associated with brain cancers (*19*). Genetic knockout of MSH3 has been shown to block somatic repeat expansion in Hdh^Q111^ mice (*13, 18, 20, 21*). Exploration of pharmaceutical approaches that selectively lower MSH3 expression in brain is warranted.

Short interfering RNA (siRNA) is a powerful therapeutic tool for sequence-specific silencing of target genes (*22, 23*). Whereas siRNA sequence defines the gene target, the scaffold (i.e., pattern of chemical modifications) of an siRNA dictates stability and delivery *in vivo* (*24-26*) Thus, once the scaffold of an siRNA has been optimized for delivery to a target tissue, any gene with a known sequence in that tissue can be targeted by changing the siRNA sequence (*27*). This programmability shortens discovery pipelines and quickens progression of compounds towards the clinic. Indeed, after establishing an siRNA architecture for delivery to the liver, four siRNA drugs were rapidly developed and approved by the FDA for treatment of liver-related conditions, with many more in late-stage clinical trials (*23, 28*). We recently developed an siRNA scaffold for delivery to the central nervous system (CNS), termed di-valent siRNA. By slowing clearance from cerebrospinal fluid and enhancing uptake into cells (*29*), di-valent siRNA support broad distribution and potent modulation of target gene expression in mouse and non-human primate (NHP) brain for up to six months after a single injection (*30*). The placement of cerebrospinal fluid (CSF) infusion (intrathecal or intracerebroventricular) has no significant impact on di-valent siRNA distribution in the large brains, confirming clinical translatability (*31*). di-valent siRNA could allow for therapeutic modulation of MSH3 expression in CNS, so long as a potent, fully modified siRNA sequence targeting *MSH3* can be identified (*30*).

Here, we identify fully chemically stabilized siRNAs targeting human, NHP and mouse *Msh3*, and show that di-valent siRNA silencing of MSH3 results in complete blockage of somatic repeat expansion over two months in two HD mouse models. Taken together, these results provide an important first step towards developing a new therapeutic paradigm for HD patients.

## RESULTS

### Identification of potent fully chemically modified siRNA sequences that silence *Msh3* mRNA in human, mouse, and non-human primate cells *in vitro*

To identify therapeutic leads for MSH3 silencing, we performed an extensive *in vitro* screen of chemically modified siRNA sequences. The panel of fully chemically stabilized siRNAs was designed using a modified siRNA efficacy prediction algorithm (*32*). In brief, the compounds were scored based on specificity, seed complement frequency, local structure, thermodynamic bias, G:C content, and positional base preferences (*32*).

From this algorithm, 48 siRNA sequences with human homology were selected. To simplify the *in vivo* validation and preclinical development path, we selected 12 additional sequences with cross-homology between mice and humans. All compounds were synthesized in an entirely modified asymmetric scaffold (*29*) with an optimized 2’-O-Methyl RNA/2’-Fluoro RNA pattern, and all terminal backbones were phosphorothioated (Fig. 1A). These chemical modifications have been shown to increase siRNA potency, stability, and duration of effect *in vivo* (*25-27, 33*). Compounds were also modified with a 3’-cholesterol conjugate on the sense strand (*24, 29*) to enable passive internalization into all cell types following addition to culture media. Sequences and chemical modification patterns for all compounds used are listed in Table S1.

**Figure 1.**
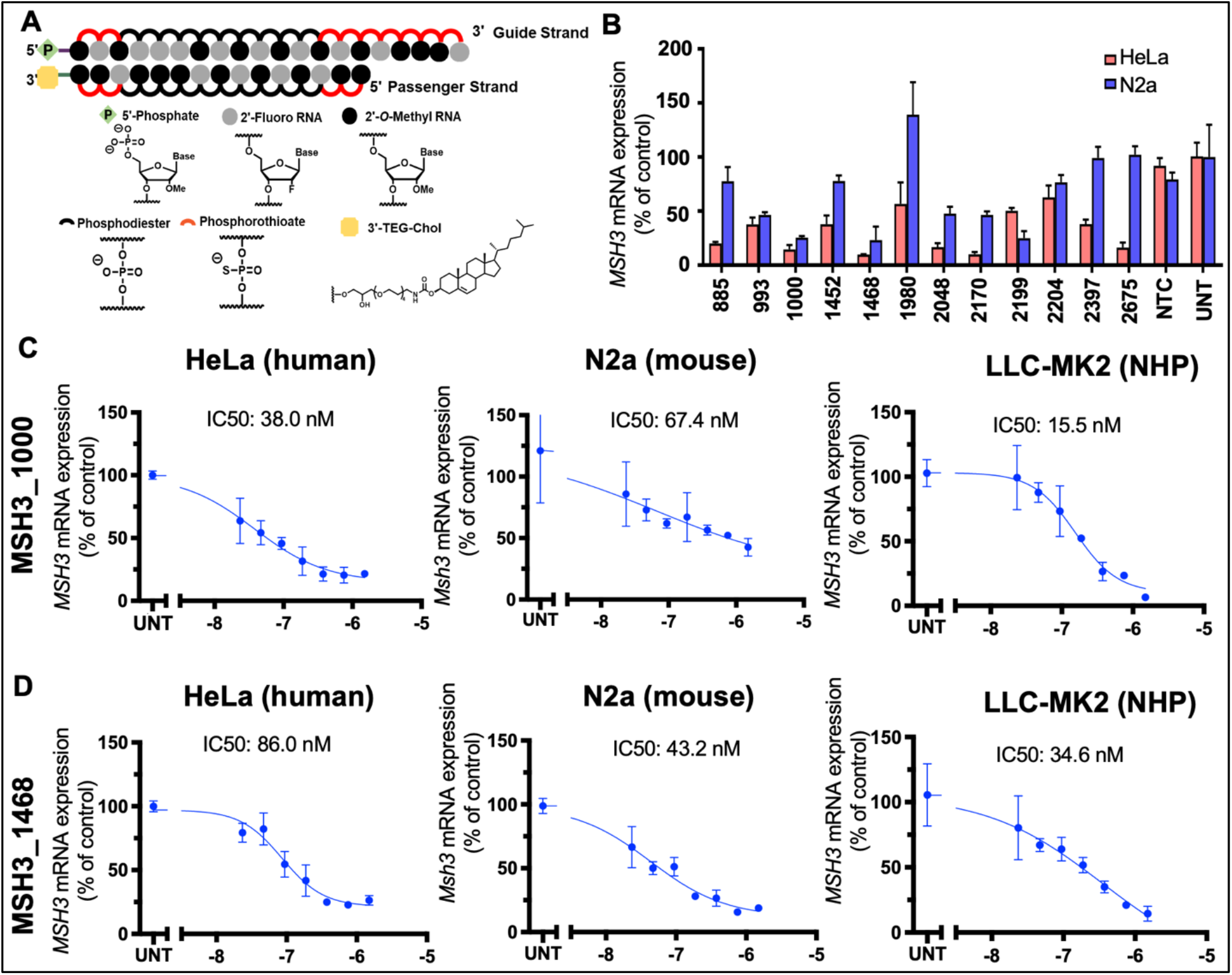
Silencing of MSH3 with fully chemically modified siRNA. **(A)** Chemical scaffold of fully modified siRNA utilized for *in vitro* screening. **(B)** *MSH3* mRNA was measured in HeLa (red) & Neuro2a (blue) cells 72 hours post-treatment with 1.5 μM siRNA or Non-Targeting Control (NTC). UNT denotes untreated controls. Dose response results for MSH3_1000 **(C)** and MSH3_1438 **(D)** in HeLa (left), N2A cells (middle), and non-human primate (NHP) LLC-MK2 cells (right). Cells treated with siRNA at concentrations shown for 72 hours. For all analyses, mRNA levels were measured using the QuantiGene Singleplex assay and calculated as percentage of untreated.

The entire siRNA panel (60 compounds) was tested in HeLa cells, and *MSH3* mRNA levels were evaluated by QuantiGene Assay at 72 hours (Fig. S1). This screen identified twelve human-targeting and six cross-reactive compounds that induce >75% silencing of *MSH3* mRNA. We next screened all twelve cross-reactive siRNAs in the mouse neuronal cell line, N2a, and identified five compounds that achieve >50% silencing of *Msh3* mRNA (Fig. 1B). The level of *Msh3* silencing in N2a cells was less pronounced than in HeLa cells. Several high efficacy compounds in HeLa cells failed to induce significant activity in N2a cells (siMSH3_1980, siMSH3_2397, siMSH3_2675). siRNA with full sequence homology can show species specific differences in maximum silencing level and overall efficacy (*34*). The observed efficacy difference may be driven by variability in the nuclear/ cytoplasmic mRNA retention and/or local changes in structural accessibility (*34, 35*).

The two cross-reactive compounds with the highest silencing efficacy in both human and mouse cells were siMSH3_1000 (86% in human and 75% in mouse) and siMSH3_1468 (90% in human and 77% in mouse). siMSH3_1000 (Fig. 1C) and siMSH3_1468 (Fig. 1D) induced dose-dependent silencing in human, mouse, and LLC-MK2 NHP cell lines (IC50s from 15-479 nM). For *in vivo* studies, siMSH3_1000 was selected given its more potent IC50 value in HeLa cells, suggesting a better pre-clinical candidate.

### Injection of di-valent siMSH3_1000 potently silences Msh3 and blocks somatic repeat expansion in striatum of Hdh^Q111^ mice

The heterozygous Hdh^Q111^ (C57BL/6J background) mouse model is a validated knock-in model of HD, in which human mutant *HTT* exon 1 is inserted in the context of the mouse *Htt* locus (*36, 37*). This model possesses a 109-111 CAG repeat tract that undergoes somatic repeat expansion within two months in the striatum (*37-39*).

To achieve robust *in vivo* efficacy and duration of effect, siMSH3_1000 and siRNA with a non-targeting control sequence (NTC) were synthesized in the di-valent scaffold (Fig. 2A)(*30*) with a 5’-Vinylphosphonate to chemically stabilize the 5’ phosphate (*33, 40*). Because silencing of *Htt* mRNA with an antisense oligonucleotide (ASO) was previously shown to block somatic repeat expansion in Hdh^Q111^ mice (*41*), we also used a previously-validated di-valent siRNA targeting *Htt* (siHTT_10150) as a control (*30, 42*). Lead or control compounds (10 nmol, or 125 μg dose) were delivered to 12-week-old Hdh^Q111^ mice (n = 6 per group) via intracerebroventricular (ICV) injection (Fig. 2B) and euthanized at 20 weeks of age to evaluate target gene silencing at the mRNA and protein level, and somatic repeat expansion.

**Figure 2.**
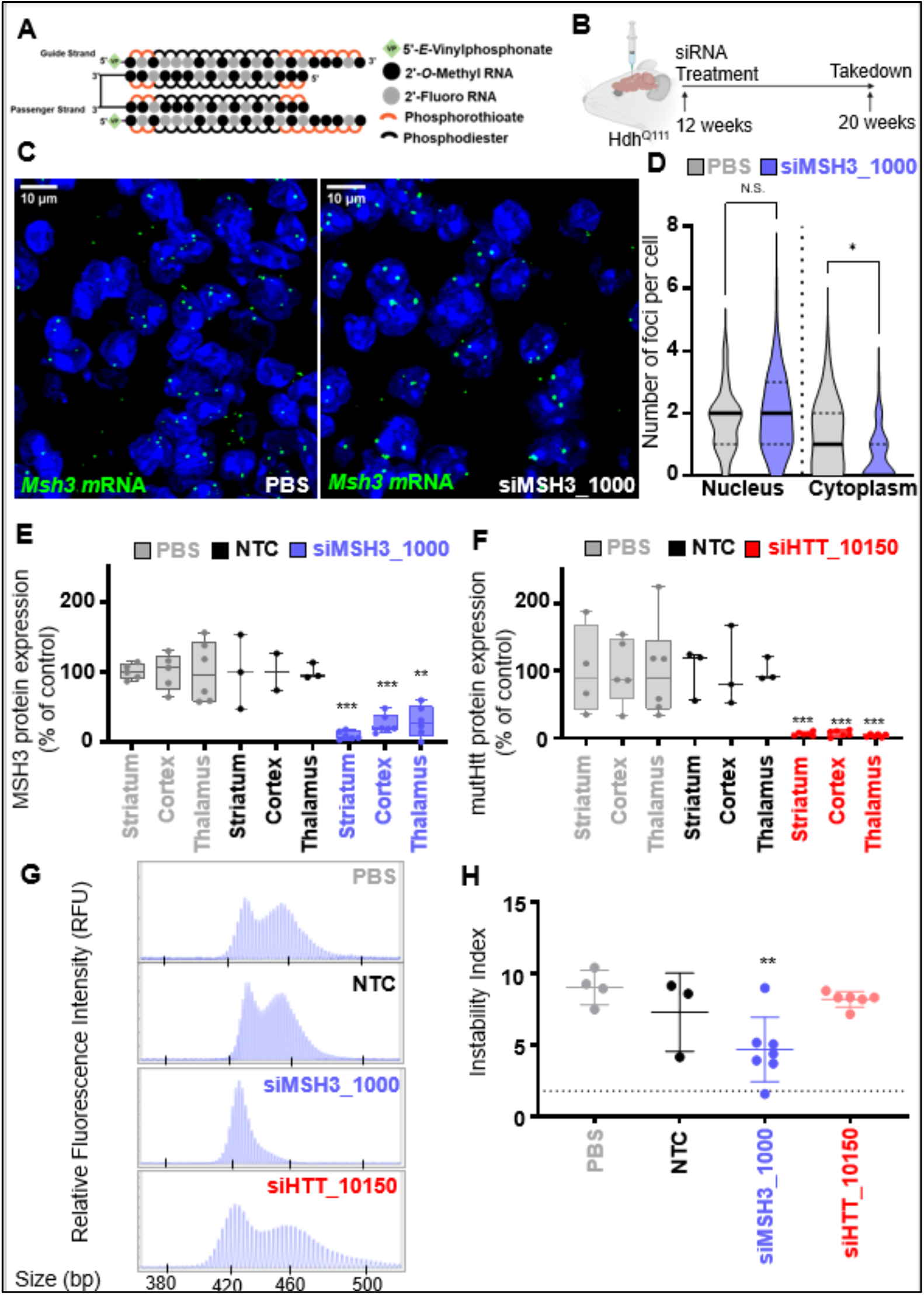
MSH3 silencing with di-siRNA blocks somatic repeat expansion in striatum of Hdh^Q111^ HD mice. **(A)** Di-valent chemically modified siRNA structure. **(B)** Experimental setup in Hdh^Q111^ mice depicts bilateral intracerebroventricular injection of PBS, di-siRNA targeting a non-targeting control (NTC), di-siHTT_10150, or di-siMSH3_1000 with 125 μg siRNA per ventricle. Mice were injected at 12 weeks old and euthanized at 20 weeks old. **(C)** *Msh3* mRNA (green) abundance and localization in striatum detected with RNAscope eight weeks post-injection following PBS (left) or siMSH3_1000 (center) treatment. Nuclei stained with DAPI (blue) **(D)** Quantification of nuclear (N) and cytoplasmic (C) foci per cell (striatum) in PBS (gray) or siMSH3_1000 (blue) treated animals. Analyzed by Kruskal-Wallis one-way ANOVA. **(E)** MSH3 protein expression and **(F)** mutant Htt (mutHTT) in striatum, cortex, and thalamus following treatment with PBS (gray), NTC (black), siHTT_10150 (red) or siMSH3_1000 (purple). Protein expression compared to NTC (one-way ANOVA with Dunnett’s multiple comparison test, *p <0.05, **p<0.01, or ***p<0.001). Each data point derives from striatum of one animal (N=4-6 animals per condition). **(G)** Representative fragment analysis of the expanded CAG locus in striatum of PBS, NTC, siMSH3_1000 and siHTT_10150 treated Hdh^Q111^ mice, eight weeks post-injection. Primers were reported in methods. **(H)** Somatic instability index calculated with a 5% signal-to-noise threshold as described in Methods. Each data point is one mouse. Instability index compared to PBS (one-way ANOVA with Dunnett’s multiple comparison test; *p <0.05, **p<0.01, or ***p<0.001).

In previous work, we showed that mutant *HTT* mRNA foci preferentially localized to the nucleus and formed clusters (*43*). Given the observed discrepancy in *Msh3* silencing level in cells of different origin, from our *in vitro* screening, it is possible that partial *Msh3* mRNA is retained in the nucleus. Therefore, to fully understand di-valent siMSH3_1000-mediated *Msh3* mRNA silencing, we used RNAscope – an advanced Fluorescence in situ Hybridization (FISH) method that allows for mRNA quantitation and visualization of nuclear/cytoplasmic localization (*44*). RNAscope uses probes that directly hybridize to the mRNA of interest and has been utilized to study mutant *HTT* mRNA clusters in HD mouse models (*43, 45*). Using RNAscope, we evaluated *Msh3* expression and intracellular localization in striatum (generated from cryosections of brain). A minimum of 150 cells were evaluated from three independent sections per animal (n=3 mice per group) as previously described (*43, 45*).

*Msh3* mRNA expression was low, with 0-4 foci detectable per cell (average of three foci), in the PBS group. As suspected, Msh3 mRNA foci were preferentially nuclear (65%) (Fig. 2C, D). In di-valent siMSH3_1000 treated brain, there was potent, near-complete silencing of cytoplasmic *Msh3* foci but no detectable impact on nuclear *Msh3* (Fig. 2D), which is consistent with siRNA-mediated mRNA silencing of Msh3 in the cytoplasm. At the protein level, di-valent siMSH3_1000 resulted in highly potent silencing (75-80% silencing, p <0.05) of Msh3 throughout the striatum, cortex, and thalamus (Fig 2E and Fig S2) compared to NTC control. This result confirms that silencing of cytoplasmic mRNA, which is the fraction that contributes to protein production, sufficiently silences protein. In di-valent siHTT_10150 treated brains, HTT protein was potently reduced (>90% silencing, p<0.01) in striatum, cortex, and thalamus (Fig. 2F and Fig S2), serving as our internal control (*30*).

Finally, we measured the effect of Msh3 and Htt silencing on somatic repeat expansion in striatum using fragmentation analysis (*46*). At 20 weeks, the PBS-treated mice had an instability index of 9.1±1.2, indicating that the two-month study window is adequate to detect somatic repeat expansion (representative traces in Fig. 2G, quantification in Fig. 2H, p<0.01). Treatment with di-valent siMSH3_1000 blocked somatic repeat expansion, keeping the instability index near baseline levels (4.7±2.2) and significantly lower than PBS controls (Fig 2H). For the NTC group, three mice died due to non-injection related issues and there is variability in the fragmentation analysis. Therefore, there is no statistical significance in the instability index in the NTC and the siMSH3_1000 treated groups. However, siHTT_10150 can be considered as a non-targeting control, since it is not targeting MSH3, but it has the same chemistry, and there is statistical differences between siMSH3_1000 (P<0.01) and siHTT_10150. Near complete silencing of Htt with di-valent siHTT_10150 had no measurable impact on somatic repeat expansion (instability index = 8.4±0.5) (Fig. 2H), contrasting previous HTT-lowering ASO results in this mouse model (*41*).

### Injection of di-valent siMSH3_1000 potently silences MSH3 and blocks somatic repeat expansion at the humanized mutant HTT locus in the BAC-CAG mouse model

To determine whether the results in the Hdh^Q111^ mouse model could be replicated in a humanized full-length mutant *HTT* context, we evaluated di-siRNA mediated modulation of MSH3 and somatic repeat expansion in the BAC-CAG HD mouse model (Fig. 3A, 3B) (*47*). BAC-CAG mice express a fully human mutant *HTT* gene with a ∼120-130 CAG repeat tract that undergoes somatic repeat expansion over two months (*47*).

**Figure 3.**
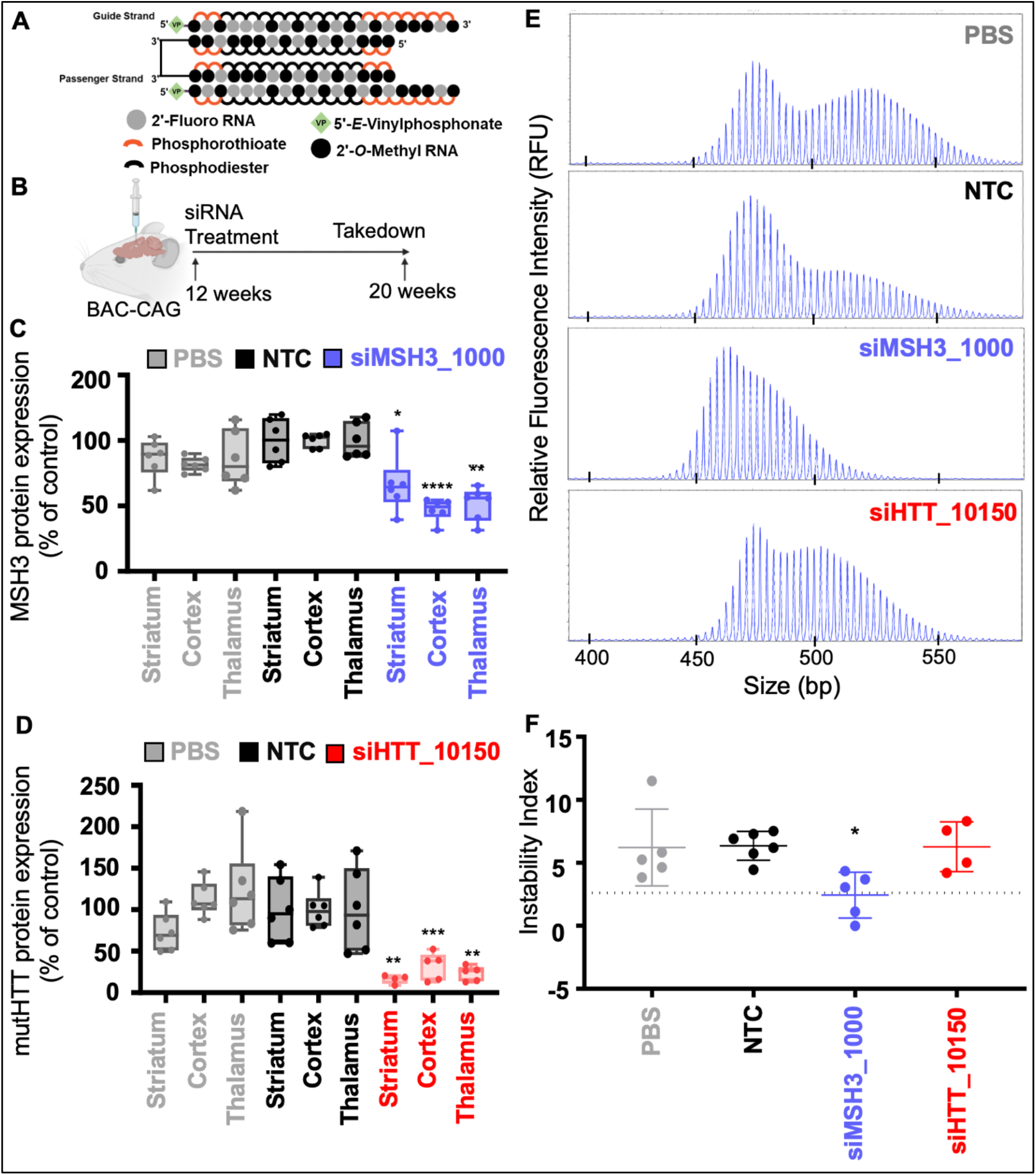
MSH3 silencing with di-siRNA blocks somatic repeat expansion in striatum of BAC-CAG HD mice. **(A)** Di-valent chemically modified siRNA structure including the chemical structure used. **(B)** BAC-CAG study plan, injecting groups at 12 weeks old: PBS, NTC, siHTT_10150, and siMSH3_1000. Mice were injected with 125 μg per ventricle of di-valent siRNA and were euthanized at 20 weeks. (C) MSH3 protein measured in PBS, NTC and siMSH3_1000 groups showing 40-50% silencing of the Msh3 protein in the striatum, cortex, and thalamus. (D) mutHTT protein expression of PBS, NTC and siHTT_10150 showing >90% silencing in the striatum, cortex and thalamus. (E) Representative fragment analysis of the expanded CAG locus in striatum of PBS, NTC, siMSH3_1000 and siHTT_10150 treated BAC-CAG mice 8 weeks post-injection. Primers reported in methods. **(F)** Somatic instability index calculated with a 5% signal-to-noise threshold as described in methods. Each data point is one mouse. Instability index compared to NTC (one-way ANOVA treatment with Dunnett’s multiple comparison test; *p <0.05, **p<0.01, ***p<0.001, ****p<0.0001).

Di-valent siMSH3_1000, siHTT_10150, PBS, or NTC (10 nmol, or 125 μg) were delivered to 12-week-old BAC-CAG mice via ICV injection and sacrificed at 20 weeks of age (Fig. 3B). Di-valent siHTT_10150 (Fig. 3C) and siMSH3_1000 (Fig. 3D) silenced HTT (80% silencing, p<0.01) and MSH3 (50% silencing, p<0.05) protein, respectively, in the striatum, cortex, and thalamus two months post-injection (Fig. S3) compared to NTC control. The level of silencing was lower than that observed in Hdh^Q111^ mice (80% vs 95% HTT silencing, 50% vs 80% MSH3 silencing).

The baseline instability index in striatum of BAC-CAG mice was 2.1±1.5. Di-valent siMSH3_1000 treated mice maintained this index (2.3±1.8), whereas NTC-treated animals had an instability index of 6.2±1.2 (representative traces in Fig 2E, quantification in Fig. 2F), suggesting that 50% silencing of MSH3 protein is sufficient to block somatic repeat expansion in the striatum. Treatment with siHTT_10150 produced no difference in the instability index (6.3 ±1.7).

### Silencing of MSH3 with MSH3_1000 or MSH3_1468 blocks somatic repeat expansion at 4 months in BAC-CAG HD mice

To determine if blocking of somatic expansion by silencing of MSH3 with divalent siRNA was robust across siRNA sequences and time points, we next used a second sequence of siRNA against MSH3 to silence MSH3 (siMSH3_1468) and measured somatic repeat expansion in the striatum at four months of treatment duration. Di-valent NTC, siMSH3_1000, siMSH3_1468 or PBS (10 nmol, or 125 μg) was delivered to BAC-CAG mice aged 12 weeks and sacrificed at 28 weeks (Fig. 4A, B). Divalent siMSH3_1000 silenced Msh3 mRNA and protein (Fig. 4C, D, S4, 70% and 65% silencing, p<0.001 and p<0.05 respectively). Di-valent siMSH3_1468 silenced Msh3 mRNA and protein (Fig. 4D, 60% and 5% silencing, p<0.001 and non-significant, respectively) compared to NTC control. The baseline somatic instability index was -0.50 ± 0.31. NTC treated striatum has an instability index of 4.14 ± 1.8. Di-valent siMSH3_1000 treated striatum had an instability index near baseline at 0.22 ± 0.47 (Fig. 4E, F, p<0.001 one-way ANOVA vs NTC), showing blocked somatic instability with 60% protein reduction. Di-valent siMSH3_1468 treated striatum had an instability index of 2.68 ± 3.4 (p<0.05 one-way ANOVA vs NTC) showing blocked somatic expansion but a wider distribution than MSH3_1000 treated tissue, suggesting a dose-dependent relationship between MSH3 silencing and blocked somatic expansion.

**Figure 4.**
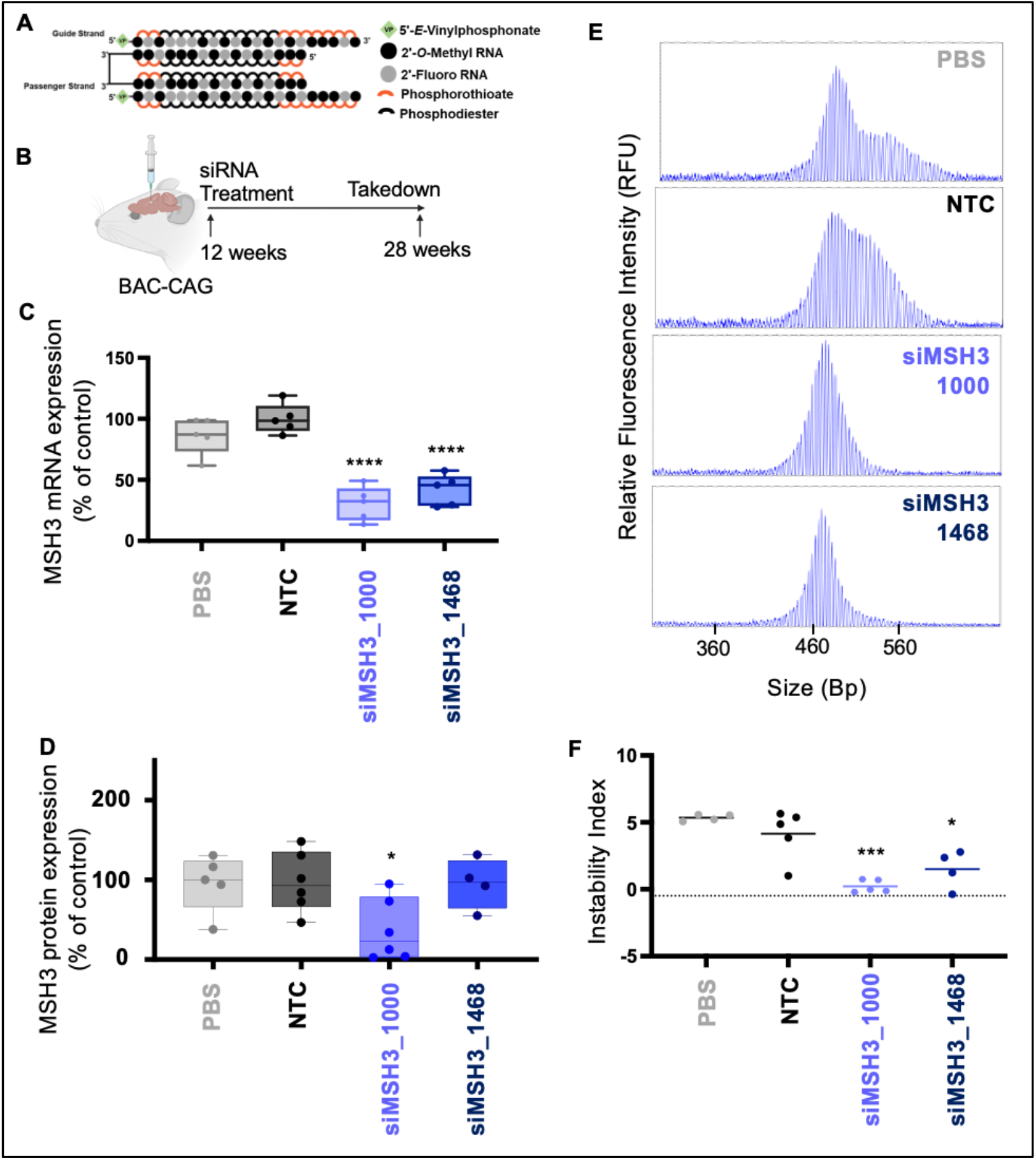
MSH3 silencing with di-valent MSH3_1000 and MSH3_1468 blocks somatic repeat expansion in BAC-CAG HD mice after 4 months treatment duration. **(A)** Di-valent chemically modified siRNA structure including the chemical structure used. **(B)** BAC-CAG study plan, injecting groups at 12 weeks old: PBS, NTC, siMSH3_1000, siMSH3_1468. Mice were injected with 125 μg per ventricle of di-valent siRNA and were euthanized at 28 weeks. **(C)** MSH3 mRNA measured in PBS, NTC, siMSH3_1000 and siMSH3_1468 groups showing 70% and 60% Msh3 mRNA silencing (respectively) in the striatum. **(D)** MSH3 protein measured in PBS, NTC, siMSH3_1000 and siMSH3_1468 groups showing 60% silencing of the Msh3 protein in the striatum with MSH3_1000 and 5% with siMSH3_1468. **(E)** Representative fragment analysis of the expanded CAG locus in striatum of PBS, NTC, siMSH3_1000 and siMSH3_1468 treated BAC-CAG mice 12 weeks post-injection. Primers reported in methods. **(F)** Somatic instability index calculated with a 5% signal-to-noise threshold as described in methods. Each data point is one mouse. Instability index compared to NTC (one-way ANOVA treatment with Dunnett’s multiple comparison test; *p <0.05, **p<0.01, ***p<0.001, ****p<0.0001).

### Silencing of MSH3 has no impact on CNS microsatellite instability

Select MMR deficiency is associated with microsatellite instability in several cancers, including colon, gastric, and endometrial cancer (*48*). To investigate whether di-valent siMSH3_1000 silencing of MSH3 alters CNS microsatellite instability, we probed three validated microsatellite loci from the Bethesda panel, which characterizes known unstable loci, that have been identified in mouse tumors: mouse big adenine tract (mBAT) 24, mBAT 26, and mBAT 64 (*49*). The lengths of each tract were measured at 4 months in di-valent PBS (N=4), NTC (N=5), siMSH3_1000 (N=5), and siMSH3_1468-treated (N=4) tissue (Fig. 5). There was no measurable difference in microsatellite instability at the mBAT 24 loci (p = 0.99), mBAT 26 loci (p = 0.89), or mBAT 64 loci (p = 0.99) across treatment groups.

**Figure 5.**
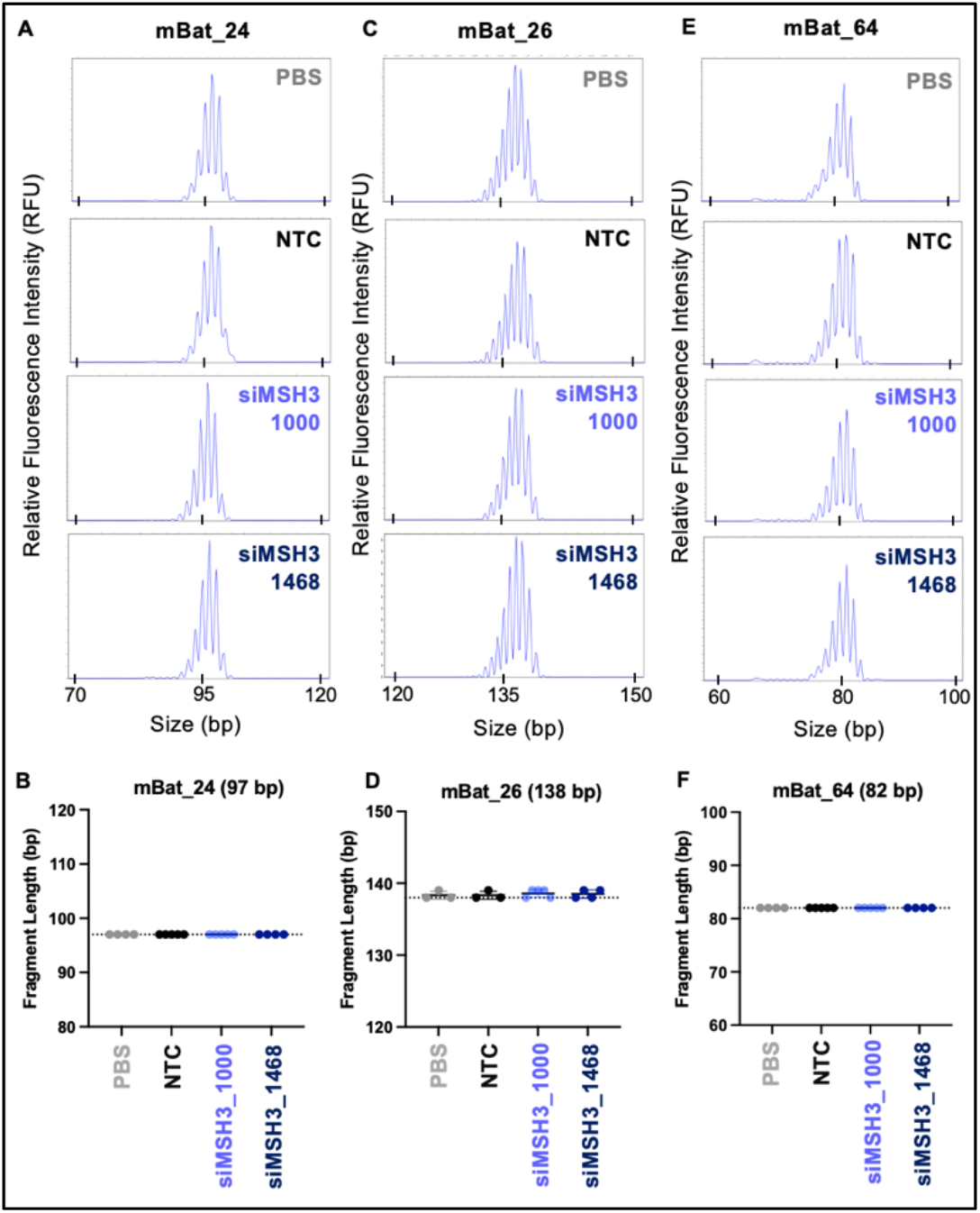
MSH3 silencing with di-valent siMSH3_1000 or si-MSH3_1468 does not affect microsatellite instability. Microsatellite instability (MSI) traces of the mBat 24 **(A)**, mBat 26 **(C)** or mBat 64 **(E)** mononucleotide repeat locus. MSI compared between PBS, NTC, siMSH3_1000 or siMSH3_1468. MSI quantitation at the **(B)** mBAT24, **(D)** mBat26, or **(F)** mBat64 loci. Traces analyzed with ThermoFischer Cloud PeakScanner. Primers reported in methods. MSI compared to NTC (one-way ANOVA).

## DISCUSSION

We demonstrate silencing MSH3 with a single dose of di-valent siRNA blocks the disease-accelerating somatic repeat expansion in mouse models of HD for up to 4 months. Somatic expansion of CAG repeats has been identified as a critical driver of HD (*13*). CAG expansion is thought to mediate its pathogenic effect through toxic downstream events at the RNA and protein level. The complexities of these downstream events have made it difficult to identify the most relevant pathogenic target for intervention (*50*). Currently, direct modulation of HTT expression is the predominant therapeutic paradigm under evaluation (*51*), but it has demonstrated limited clinical success (*50*). Targeting the potential accelerator of pathogenesis – i.e., expansion of CAG repeats –may slow, or even stop disease progression. Moreover, somatic repeat expansion is a key feature of other trinucleotide repeat disorders, including myotonic dystrophy and Friedreich’s ataxia, making this therapeutic approach applicable to any diseases associated with the somatic repeat expansion.

Although GWAS data have identified several MMR genes as both HD and somatic repeat expansion modifiers, mounting evidence suggests a strong association between lower MSH3 expression, reduced somatic repeat expansion, and slower disease onset/progression (*13, 18, 52*). Genetic knockout of MSH3 clearly shows blocking of somatic repeat expansion and thus provides independent confirmation of the critical role of MSH3 in somatic repeat expansion (*21*). Targeting MSH3 is a promising therapeutic direction for HD. To validate siRNA-mediated modulation of MSH3 *in vivo*, we chose two mouse models of HD, one with the mouse *htt* locus and the other with the human *HTT* locus (BAC-CAG), both of which undergo CAG repeat expansion. The BAC-CAG mouse is the first HD model with uninterrupted CAG repeats within the full human mutant HTT gene, which undergoes expansion (*47*). The observed difference in levels of MSH3 silencing between the mouse models is intriguing and could be due to the different backgrounds of each model, C57BL/6 (for Hdh^Q111^) versus FVB (for BAC-CAG), which might express different isoforms of MSH3 or have different polymorphisms of MSH3 (*53*). Another explanation could be that the mRNA is structured differently in the two backgrounds, making the target site less available.

We found that di-siRNA-mediated modulation of HTT had no effect on somatic repeat expansion *in vivo*. This finding contradicts an early preprint study in which antisense oligonucleotide (ASO)-mediated silencing of *Htt* reduced somatic repeat expansion (*41*). It is likely that the *Htt*-targeting ASOs reduce somatic repeat expansion by interfering with locus transcriptional rates through binding nascent nuclear targets. (*54*). By contrast, siRNA-mediated silencing of mRNA in the cytoplasm would have no impact on the somatic repeat expansion.

MSH3 silencing and blockade of somatic expansion were greater with siMSH3_1000 than siMSH3_1468 suggesting that the level of MSH3 is tightly connected to the extent of somatic expansion. The exact amount of MSH3 lowering that is required to block somatic expansion in patients is an outstanding question for further study.

The clinical utility of targeting MSH3 in HD relies heavily on the safety of lowering its expression. MSH3 selectively recognizes long DNA loops and recruits other MMR machinery for DNA repair (*55*). This highly specific role of MSH3 in MMR explains its limited association in cancers (*16, 56*), with no known relationship to CNS-derived tumors being reported (*19*). Our findings showing no effect of MSH3 lowering on tumor-associated microsatellite instability provides further evidence for the safety of this therapeutic approach. Ongoing research is required to study, potential long term toxicity associated with silencing of MSH3 and investigating the impact on neuronal damage markers.

Additionally, the pharmacokinetics and pharmacodynamics (PK/PD) of Di-siRNA makes it a safe and effective drug modality for MSH3 modulation, where majority of the injected dose is retained in the CNS. Di-valent siRNAs, when administered by CSF infusion, achieve 30-40% retention of the injected dose in the CNS (*31*) (*57*), where it broadly distributes (*30*) –to brain structures highly affected in HD –and can silence a gene target for up to six months. Importantly, di-valent siRNAs show limited accumulation in liver and kidney and no detectable presence in the colon, a tissue in which MSH3 silencing is associated with cancer (*58, 59*).

While the MSH3 is an obvious top target to explore for modulation of somatic repeat expansion, other genes involved in the MMR pathway might be of interest. For example, genetic knockout of MLH1 and MLH3 have been shown to impact somatic repeat expansion in HD mouse models (*13, 36, 60, 61*). The inherent sequence specificity of siRNAs and their duration of effect provide a powerful therapeutic paradigm for treating neurodegenerative disorders. The discovery and development of di-valent (*30*) and lipophilic siRNA (*62, 63*) has opened siRNA drugs to CNS indications; several compounds are now in clinical or late preclinical development (*58*).

While somatic repeat expansion contributes to HD, silencing of the expanded HTT protein might still be necessary at late stages of disease (*64*). Depending on disease progression at time of treatment, simultaneous modulation of somatic repeat expansion and mutant HTT expression might be required. One advantage of siRNA is the ability to make “cocktails” of siRNA that target multiple genes simultaneously. Thus, ongoing studies are investigating the effects of combinatorial HTT and MSH3 silencing on disease progression in HD mouse models.

In conclusion, somatic repeat expansion has been clearly identified as a possible therapeutic target to stop progression of HD. We developed a potent, fully chemically stabilized, cross species di-valent siRNA that lowered MSH3 expression and block CAG repeat expansion *in vivo*. This approach represents a new therapeutic paradigm for HD as well as other repeat diseases where somatic expansion contributes.

## MATERIALS AND METHODS

### Oligonucleotide synthesis, Quality Control, and siRNA preparation for screening

Oligonucleotides were synthesized by phosphoramidite solid-phase synthesis on a Dr Oligo 48 (Biolytic, Fremont, CA) using 2’-Fluoro RNA or 2’-O-Methyl RNA phosphoramidites with standard protecting groups purchased from ChemGenes, Wilmington, MA. Non-conjugated oligonucleotides were synthesized on a 500 Å UnyLinker support (ChemGenes). Cholesterol conjugated oligonucleotides were synthesized on a 500 Å tetraethylene glycol cholesterol support (ChemGenes). Phosphoramidites were prepared at 0.1 M in anhydrous acetonitrile (ACN), except for 2’-O-methyl-uridine phosphoramidite, which was dissolved in anhydrous ACN containing 15% anhydrous *N,N*-Dimethylformamide (DMF). To activate the phosphoramidites, 5-(Benzylthio)-1*H*-tetrazole (BTT) (0.25 M) in anhydrous ACN was used and the coupling time was four minutes. Capping of unreacted sites was performed using CAP A (20% 1-Methyl-*1H*-imidazole in ACN) and CAP B (30% 2,6-Lutidine and 20% acetic anhydride in ACN). To oxidize the phosphite (P III) centre to the phosphate (P V) centre, 0.05 M iodine in pyridine-water (9:1, v/v, AIC) was added for four minutes. To sulfurize the phosphite centers, a 0.1 M solution of 3-[(dimethylaminomethylene)amino]-3*H*-1,2,4-dithiazole-5-thione (DDTT) in pyridine (ChemGenes) was added for four minutes. For detrytilation reactions, 3% trichloroacetic acid in dichloromethane (AIC) was utilized. Post synthesis, the columns were washed with a solution of 10% *N,N*-Ethylethaneamine (DEA) in anhydrous ACN.

For deprotection of oligonucleotides, methylamine gas (purchased from Airgas) was used for one hour at room temperature in a gas chamber. The oligonucleotides were placed in a vacuum desiccator to help remove any remaining methylamine gas for 20 minutes. The columns were washed with 0.1 M sodium acetate 80% Ethanol in water (five times), followed by 85% ethanol in water (five times) to precipitate oligonucleotides on the support. The ethanol was removed by heating for five minutes and placing columns in a vacuum desiccator for 20 minutes. The final oligonucleotides were eluted with water. Identity of the oligonucleotides was confirmed via LC-MS.

Oligonucleotides were quantitated using a TECAN SPARK system by measuring the absorbance at 260 nm (A_260_) of a 1:40 dilution with water. The concentrations were calculated using Beers law. The complementary antisense and sense strand were then combined to make a final siRNA concentration of 100 μM in water. Finally, this solution was heated to 95°C for five minutes and allowed to cool down to room temperature over an hour to anneal the two strands.

### Synthesis of oligonucleotides for *in vivo* injections

Oligonucleotides for *in vivo* studies were synthesized on a MerMade 12 automated oligonucleotide synthesizer. Di-valent oligonucleotides were synthesized on a solid support that has been previously published (*30*). For oligonucleotides that required 5’-(*E*)-Vinyl tetra phosphonate (pivaloyloxymethyl), 2’-*O*-methyl-uridine 3’-CE phosphoramidite (VP) was purchased from Hongene. di-valent oligonucleotides were synthesized on a modified solid support (*30*). The synthesis cycle procedure is the same as previously discussed for screening. For deprotection of VP-containing oligonucleotides, 3% DEA in Ammonia hydroxide was used and heated at 30°C for 20 hours. di-valent oligonucleotides were deprotected using an aqueous Ammonia hydroxide (28-30% in water): Methylamine solution (40% in water) (1:1, v/v) at RT for two hours. The solutions were evaporated to dryness. Crude material was dissolved in water and filtered to remove CPG. The crude material was purified on an Agilent 1200 Prep HPLC system using Ion Exchange. After purification, oligonucleotides were desalted on an AKTA FPLC with Sephadex columns. Quantitation of oligonucleotides were performed on a Nanodrop system. To anneal siRNA, the antisense and sense were added together and heated to 95°C for five minutes before being allowed to cool to room temperature slowly.

### Cell Culture

HeLa (#CCL-2), Neuro-2a (#CCL-131), and LLC-MK2 (#CCL-7) cells were purchased from the American Type Culture Collection (ATCC). HeLa and LLC-MK2 cells were maintained with Dulbecco’s Modified Eagle’s Medium (DMEM) (Cellgro, #10-013CV) with 10% fetal bovine serum (FBS) (Gibco, #26140) and Neuro-2a cells were maintained with EMEM (ATCC, #30-2003) with 10% FBS. Both cell lines were cultured at 37°C and 5% CO_2_. Cells were split once confluent and were limited to 20 passages.

### Screening and dose responses

For screening, siRNA were diluted in OptiMeM (Carlsbad, CA; 31985-088) to double their final concentration, then 50 μl of this dilution was placed in triplicate on a 96-well plate. Cells were trypsinized, centrifuged, and resuspended in 6% media. This suspension was counted and diluted in 6% FBS media so that there were 8,000-12,000 cells per 50 μL. These cells were added to the siRNA in the 96-well plate. These additions resulted in a final FBS amount of 3% and 1.5 μM concentration of siRNA. Plates were incubated for 72 hours at 5% CO_2_ and 37°C.

For 7-point dose response studies, siRNA was diluted in OptiMeM then serially diluted (2x dilution) and plated in triplicate on 96-well plates. For the addition of cells, the same procedure outlined for screening was followed.

### Animal Experiments

All animal care was within accordance with institutional guidelines. Animal experiments were approved by the UMass Chan Medical School IACUC (protocol 202000010). Hdh^Q111^ mice were provided from Jackson Laboratory, Bar Harbor, Maine, USA at 6-8 weeks of age. At 12 weeks of age, the mice were bilaterally ICV injected with 10 μL of siRNA (5 μL per ventricle) in PBS, at a rate of 500 nl min^-1^. The coordinates from Bregma are: -0.2 mm AP, ± 1.0 mm mediolateral and -2.5 mm dorsoventral. Mice were anesthetized throughout the procedure using 1.2% Avertin or isoflurane. After two months, mice were euthanized. Mice were perfused with PBS and half brains were frozen in OCT for RNAscope. For protein analysis, 1.5 mm x 1.5 mm punches were flash frozen; for mRNA analysis, punches were placed in RNA later for 24 hrs at 4°C.

### Western blot analysis

Frozen tissue punches from striatum, medial cortex, posterior cortex, and thalamus were homogenized on ice in 75 μl 10 mM HEPES pH 7.2, 250 mM sucrose, 1 mM EDTA + protease inhibitor tablet (Roche, complete, EDTA-free) + 1 mM NaF + 1 mM Na_3_VO_4_, and sonicated for 10 seconds. Protein concentration was determined using the Bradford method (BioRad). Equal amounts of protein (10 mg) were separated by SDS-PAGE and analyzed by western blot using antibodies to Huntingtin (1:2000, Ab1, aa1-17,(*65*) MSH3 (1:500, Santa Cruz) and b-tubulin (1:5000, Sigma), and GAPDH (1:10,000, Millipore) as previously described (*66*). Bands were visualized with SuperSignal West Pico PLUS Chemiluminescence substrate (Pierce) and images were obtained with a CCD camera (AlphaInnotech). Pixel intensity quantification was performed using ImageJ software (NIH) by manually circling each band and multiplying the area by the average intensity to obtain the total intensity for each band and normalizing the signal to the tubulin or GAPDH loading control.

### RNAscope

*MSH3* RNA was visualized via *in situ* RNA hybridization using the multifluorescent RNAScope^tm^ assay kit (Advanced Cell Diagnostics INC). Probes for *MSH3* (Cat No. 428921), were applied to fresh-frozen 20um thick brain tissue sections in accordance with manufacturer’s instructions, and resulting slides were imaged using a Leica SP8 confocal microscope. For controls, a negative control probe set (Cat No. 320871) and a positive control probeset (Cat No. 320881) were used. Quantitative analyses for RNA abundance and nuclear: cytoplasmic localization were developed in-house using FIJI (NIH).

### Fragmentation analysis

Genomic DNA was extracted from ∼10 mg punches from selected brain regions using the solid tissue protocol from the IBI gMax Mini Kit (IBI cat# IB47218). DNA concentrations were determined using the Qubit Flex Fluorometer. The CAG repeat region of the Htt gene was amplified in 80 μL PCR reactions using forward primer CAG1 (ATGAAGGCCTTCGAGTCCCTCAAGTCCTTC, 6FAM-labeled) and reverse primer Hu3 (GGCGGCTGAGGAAGCTGAGGA). Each 80 μL PCR reaction consisted of 8 μL AmpliTaq buffer, 8 μL DMSO, 4 μL BSA (20 mg/mL), 8 μL GC enhancer, 3.2 μL 25 mM MgCl_2_, 8 μL 2 mM dNTPs, 6.4 μL 10 μM forward primer, 6.4 μL 10 mM reverse primer, 19.2 μL H_2_O, and 0.8 μL AmpliTaq 360 Taq Polymerase. Thermocycling conditions were as follows: An initial denaturation of 1 minute 30 seconds at 94 °C, then 35 cycles of 30 seconds at 94 °C, 30 seconds at 63.5 °C, and 1 minute 30 seconds at 72 °C, followed by a final annealing step of 10 minutes at 72 °C. The 80 μL PCR product was concentrated down to 20 μL using the GeneJET PCR Purification kit (Thermo Fisher cat# K0702). PCR products were eluted into 20 μL of water. Concentrated PCR products were sent to GeneWiz for fragment analysis. Traces were visualized using Peak Scanner 2 software, and expansion indices were calculated using a custom R script based on somatic instability index calculations (*46*).

Statistics: When comparing two or more groups, one-way ANOVA vs control (NTC, PBS) with Dunnet’s post-hoc analysis was used. When comparing two groups, a Student’s t-test was used. * p ≤ 0.05; ** p ≤ 0.01; *** p ≤ 0.001; **** p ≤ 0.000.

## Acknowledgments

Authors acknowledge all Khvorova and Aronin lab members who kindly reviewed the manuscript and provided editorial suggestions.

## Funding

MD and NA are funded under NIH U01 NS114098. MD thanks CHDI-6367 and the Dake Family fund as funding sources. NA is funded under NIH R01 NS106245. NA and AK thank the CHDI Foundation. AK is funded under NIH R35 GM131839 and S10 OD020012.

## Author contributions

Conceptualization: CF, AK, JB

Visualization: DO, JB, AK, NA, MD

Methodology: DO, JB, CF, AK

Investigation: DO, JB, AK, CF, AS, ES, CM, EM, JB, SL, DME, ZK, VH, KM

Funding acquisition: AK, NA, MD, JC,

Project administration: AK, NA, MD, JC,

Supervision: AK, NA, MD, JC,

Writing – original draft: DO, JB, AK,

Writing – review & editing: DO, JB, AK, NA, MD

## Competing interests

NA and AK are co-founders of Atalanta, on the Scientific Advisory Board of Atalanta, Huntington’s Disease Society for America, and Biogen, and has licensing to Spark Therapeutics. AK has a financial interest in Atalanta and is on the Atalanta Scientific Advisory Board. The authors have patents on the MSH3 as a therapeutic target and methodology described in the paper.

## Data and materials availability

All data are available in the main text or the supplementary materials.

