## Supplementary figures and images for "Di-valent siRNA Mediated Silencing of MSH3 Blocks Somatic Repeat Expansion in Mouse Models of Huntington’s Disease"

### Supporting Information

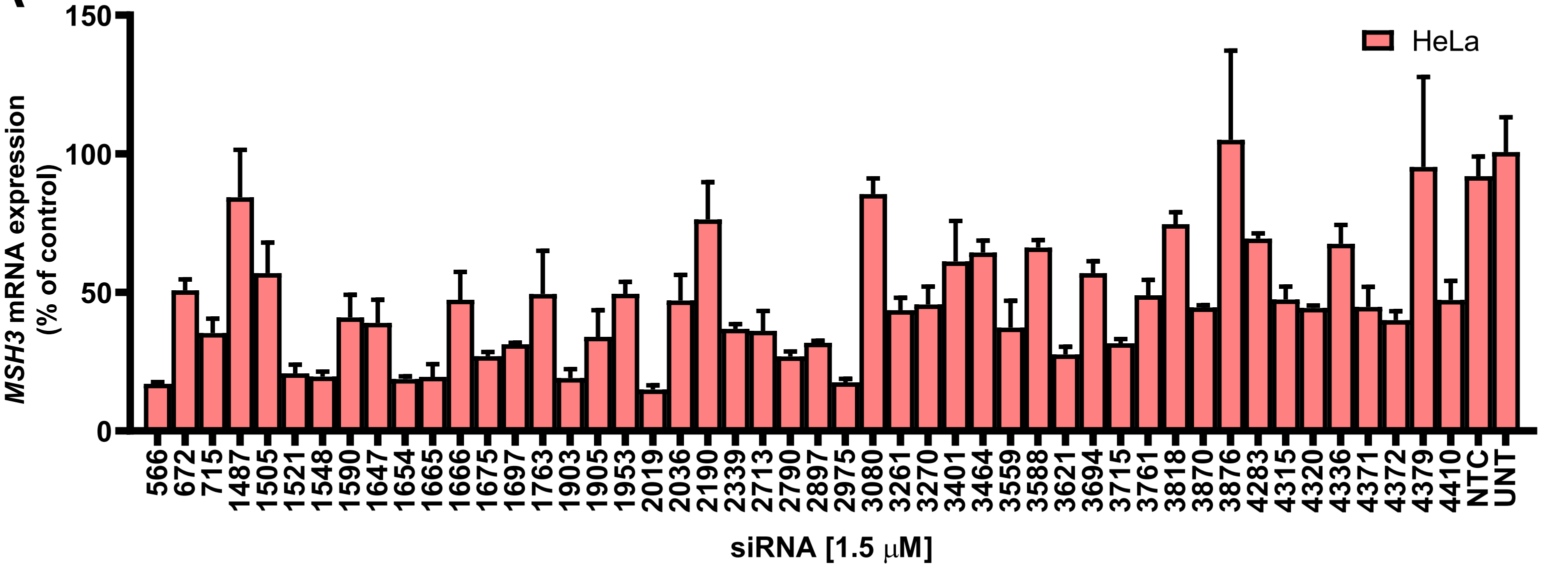

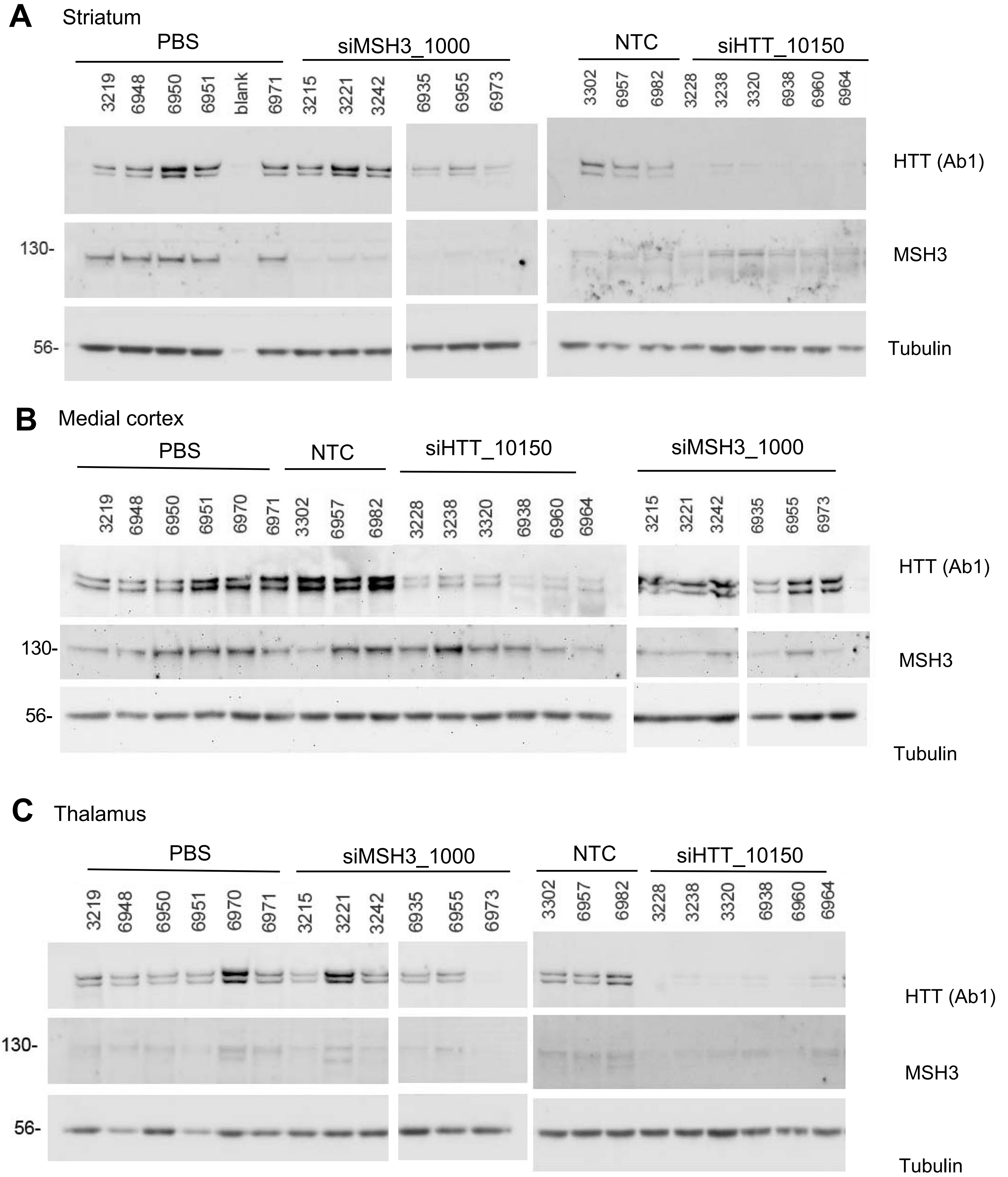

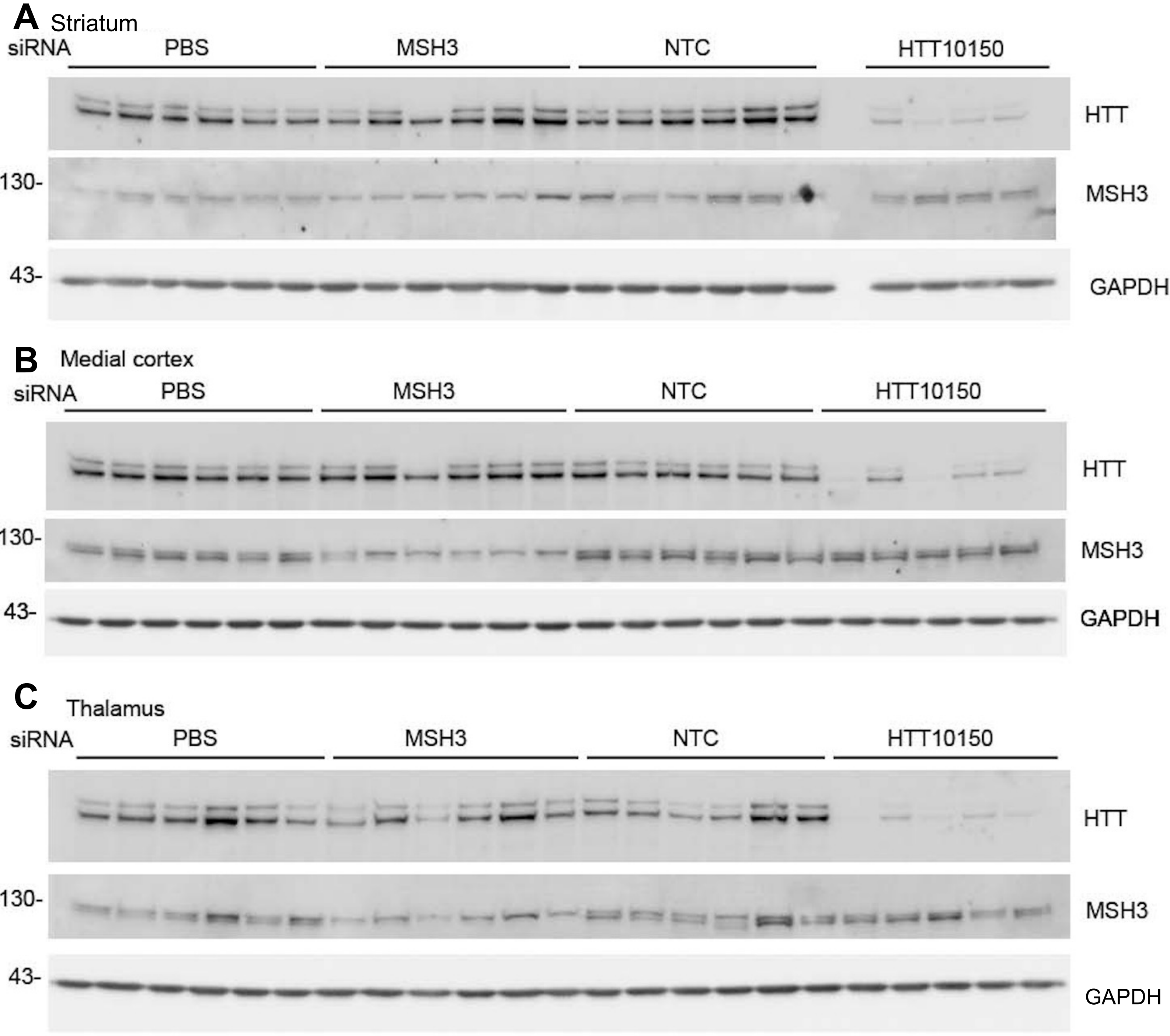

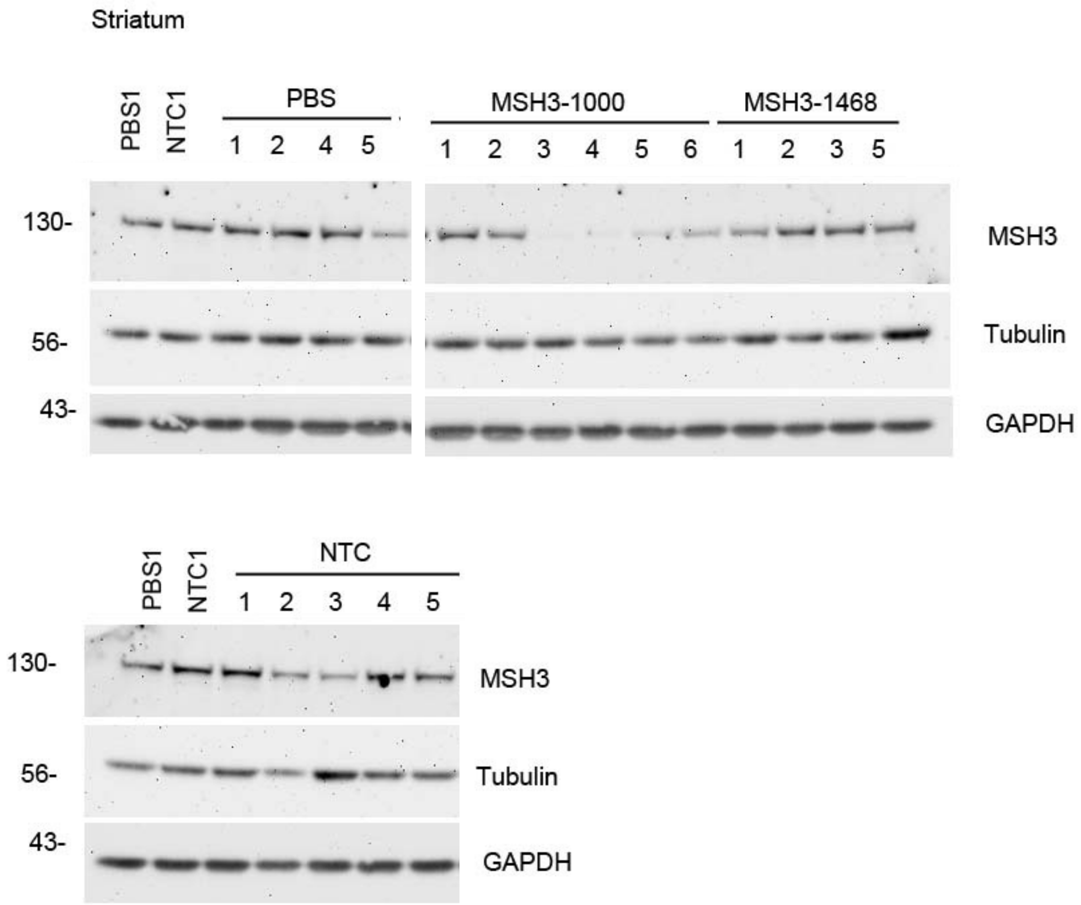
